# Genome-wide analysis of *Escherichia coli* fitness determinants in a cross-feeding mutualism with *Rhodopseudomonas palustris*

**DOI:** 10.1101/2020.02.20.958660

**Authors:** Breah LaSarre, Adam M. Deutschbauer, Crystal E. Love, James B. McKinlay

## Abstract

Microbial interactions abound in natural ecosystems and shape community structure and function. Substantial attention has been given to cataloging mechanisms by which microbes interact, but there is a limited understanding of the genetic landscapes that promote or hinder microbial interactions. We previously developed a mutualistic coculture pairing *Escherichia coli* and *Rhodopseudomonas palustris*, wherein *E. coli* provides carbon to *R. palustris* in the form of glucose fermentation products and *R. palustris* fixes N_2_ gas and provides nitrogen to *E. coli* in the form of NH_4_^+^. The stable coexistence and reproducible trends exhibited by this coculture make it ideal for interrogating the genetic underpinnings of a cross-feeding mutualism. Here, we used random barcode transposon sequencing (RB-TnSeq) to conduct a genome-wide search for *E. coli* genes that influence fitness during cooperative growth with *R. palustris*. RB-TnSeq revealed hundreds of genes that increased or decreased *E. coli* fitness in a mutualism-dependent manner. Some identified genes were involved in nitrogen sensing and assimilation, as expected given the coculture design. The other identified genes were involved in diverse cellular processes, including energy production and cell wall and membrane biogenesis. Additionally, we discovered unexpected purine cross-feeding from *R. palustris* to *E. coli*, with coculture rescuing growth of an *E. coli* purine auxotroph. Our data provide insight into the genes and gene networks that can influence a cross-feeding mutualism and underscore that microbial interactions are not necessarily predictable *a priori*.

**IMPORTANCE:** Microbial communities impact life on earth in profound ways, including driving global nutrient cycles and influencing human health and disease. These community functions depend on the interactions that resident microbes have with the environment and each other. Thus, identifying genes that influence these interactions will aid the management of natural communities and the use of microbial consortia as biotechnology. Here, we identified genes that influenced *Escherichia coli* fitness during cooperative growth with a mutualistic partner, *Rhodospeudomonas palustris*. Although this mutualism centers on the bidirectional exchange of essential carbon and nitrogen, *E. coli* fitness was positively and negatively affected by genes involved in diverse cellular processes. Furthermore, we discovered an unexpected purine cross-feeding interaction. These results contribute knowledge on the genetic foundation of a microbial cross-feeding interaction and highlight that unanticipated interactions can occur even within engineered microbial communities.

## INTRODUCTION

Within every ecosystem, microbial cells interact with both the environment and each other, and these interactions shape ecosystem functions (62). Microbial interactions are diverse and include nutrient competition and cross-feeding (55), cell-cell signaling (1), adhesion (53), and chemical warfare (27). Unfortunately, the identification of interaction mechanisms has far outpaced our understanding of the genetic and physiological bases of microbial interactions.

Much knowledge on microbial interactions has come from synthetic communities (i.e., cocultures) that offer simplicity and tractability over natural communities (34, 62). Our lab previously developed an anaerobic coculture pairing fermentative *Escherichia coli* with the N_2_-fixing, photoheterotroph *Rhodopseudomonas palustris* (31). In this coculture, the sole carbon source is glucose, which *R. palustris* cannot utilize, and the sole nitrogen source is N_2_ gas, which *E. coli* cannot utilize. Consequently, each species depends on the other for an essential nutrient: *R. palustris* gets carbon in the form of *E. coli* fermentation products, and *E.* coli gets nitrogen in the form of NH_4_^+^ that is excreted by an engineered *R. palustris* strain (Nx) (**Fig. 1A**). The stable coexistence and reproducible trends (31, 38, 39) make this coculture ideal for investigating the genetic and physiological characteristics that influence a cross-feeding mutualism.

**Fig. 1.**
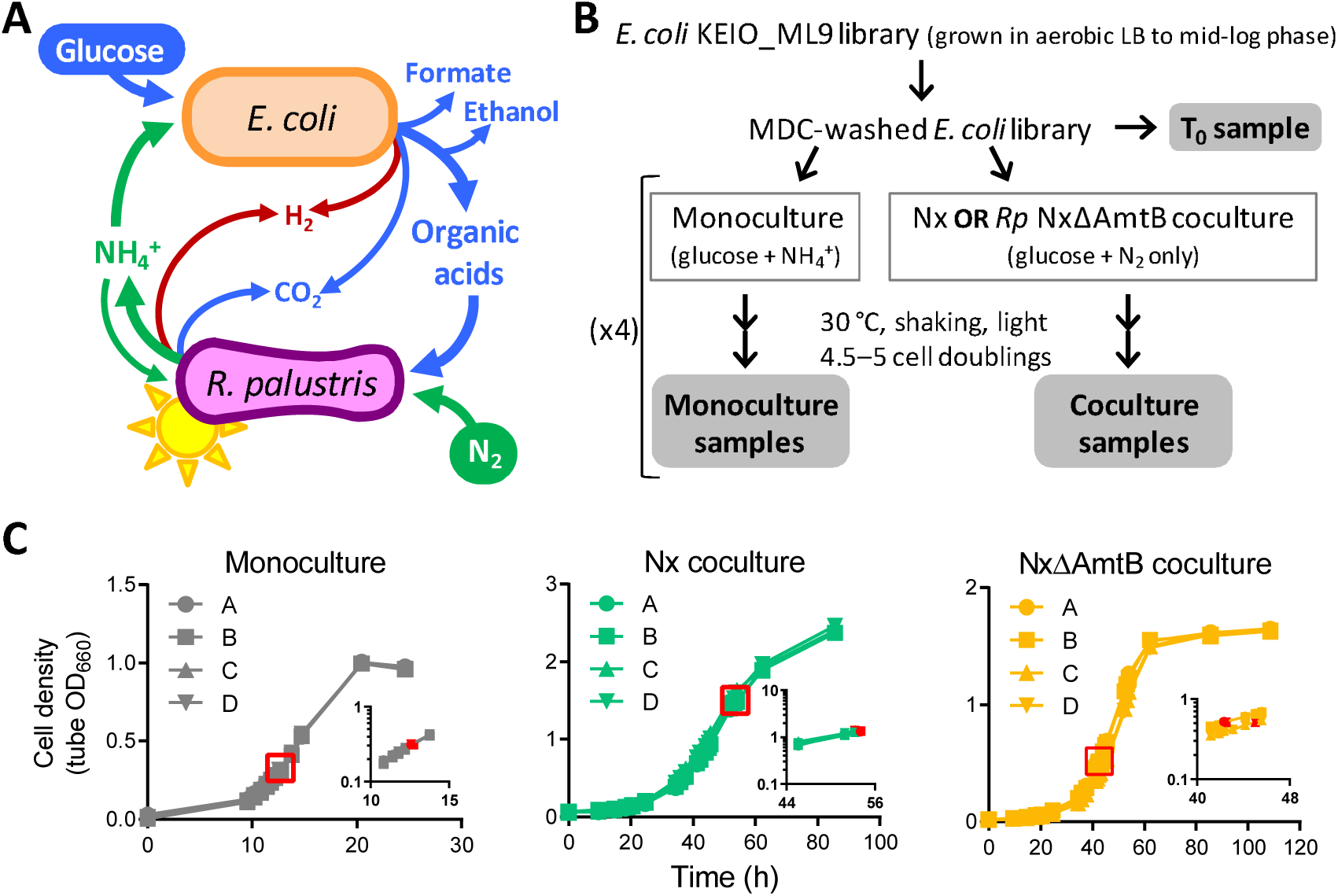
Overview of RB-TnSeq screen for *E. coli* genes that influence fitness during mutualistic growth with *R. palustris*. (A) Mutualistic growth between *E. coli* (*Ec*) and *R. palustris* (*Rp*) requires carbon transfer, in the form of glucose fermentation products, from *E. coli* to *R. palustris* and nitrogen transfer, in the form of NH_4_^+^, from *R. palustris* to *E. coli*. (B) Experimental design. There was one T_0_ sample and 4 biological replicates for all mono- and cocultures. Gray bubbles indicate samples that were used for BarSeq. (C) Quadruplicate growth curves of KEIO_ML9a monocultures or cocultures with *R. palustris* Nx or *R. palustris* NxΔAmtB. Red boxes indicate sampling points. Insets (X, time; Y, cell density) show portions of the same growth curves on a log scale. For cocultures with *R. palustris* Nx, samples were harvested at densities above the linear range of the spectrophotometer; as such, the inset graph reports cell densities measured in cuvettes with samples diluted to values within the linear range.

Recently, our lab used transcriptome sequencing (RNA-Seq) and proteomics to interrogate how cross-feeding impacts the physiology of each species (37). This study revealed that the *E. coli* NtrC-mediated nitrogen starvation response (NSR) is crucial for fitness in coculture (37). However, RNA-Seq does not necessarily identify genes that confer fitness under the tested conditions. Indeed, several highly upregulated genes did not contribute to *E. coli* fitness in coculture (37). One approach for directly identifying fitness determinants is transposon sequencing (Tn-Seq), which screens for the fitness of millions of individual transposon mutants simultaneously using next-generation sequencing (58). Random barcode Tn-Seq (RB-TnSeq) uses random barcodes to uniquely identify each Tn insertion site, thereby reducing the sample preparation needed for fitness analysis (61). RB-TnSeq has been successfully used to identify genetic fitness determinants in numerous bacteria under diverse conditions (21, 22, 50, 51), including in multispecies communities (43).

Here, we used RB-TnSeq to identify *E. coli* genes that affected fitness during cooperative growth with *R. palustris*. We identified numerous genes that had negligible impact on *E. coli* fitness in monoculture but promoted or reduced *E. coli* fitness in coculture. These mutualism-dependent fitness determinants were involved in diverse physiological processes ranging from amino acid metabolism to signal transduction to membrane biogenesis. Unexpectedly, we also identified *E. coli* genes that were important for monoculture fitness but dispensable in coculture, including genes for *de novo* purine synthesis. These results revealed an unprompted cross-feeding interaction that was not predictable *a priori*. Overall, our study provides insight into *E. coli-R. palustris* interactions and lays the groundwork for understanding the genetic underpinnings of a cross-feeding mutualism. Additionally, our results highlight that microbes can interact in unanticipated and potentially covert manners, even in engineered communities.

## RESULTS AND DISCUSSION

### Experimental setup of RB-TnSeq screen for *E. coli* fitness determinants in coculture

Previously, *Wetmore* et al. (61) generated and characterized an *E. coli* RB-TnSeq library in the K12 parent strain of the KEIO deletion collection, BW25113 (4). This library, named KEIO_ML9, comprises 152,018 uniquely bar-coded mutants covering 3,728 of the 4,146 protein-coding genes (61). To identify *E. coli* genes contributing to fitness in coculture with *R. palustris*, we grew the KEIO_ML9 library under three conditions (**Fig. 1B**): (i) “Nx coculture”, that is coculture with *R. palustris* Nx (CGA4005), a strain that excretes NH_4_^+^ at a arbitrarily defined level of 1X (31); (ii) “NxΔAmtB coculture”, that is coculture with *R. palustris* NxΔAmtB (CGA4021), a strain that excretes 3X NH_4_^+^ (31); and (iii) “monoculture”, that is monoculture using the same anaerobic medium used for cocultures but supplemented with 15 mM NH_4_Cl to enable *E. coli* growth in the absence of *R. palustris*. Monoculture was used to identify genes that affected *E. coli* fitness due to the anaerobic, minimal-medium environment. We examined *E. coli* fitness in both Nx and NxΔAmtB cocultures because *E. coli* growth is differentially constrained in these two cocultures. Specifically, the 1X NH_4_^+^ cross-feeding level in Nx coculture constrains *E. coli* growth such that it is coupled to that of *R. palustris*, with both species exhibiting a doubling time of approximately 10 h (31); in NxΔAmtB cocultures, the 3X level of NH_4_^+^ cross-feeding allows *E. coli* to grow faster than *R. palustris*, although *E. coli* still experiences a degree of nitrogen limitation and exhibits slower growth than in monocultures with excess NH_4_^+^ (31). We reasoned that genes influencing the *E. coli*-*R. palustris* interaction would exhibit fitness effects of different magnitudes in these two cocultures, with effects being more pronounced in Nx cocultures.

To enable accurate and informative comparisons between conditions, we took care in how we inoculated, incubated, and harvested the cultures. First, all cultures were inoculated using minimal-medium-washed *E. coli* cells from a single KEIO_ML9 culture grown to mid-log phase in aerobic LB with kanamycin (**Fig. 1B**). Aerobic LB served as a ‘non-selective’ condition to preserve library diversity (61). However, it was imperative to wash the cells using minimal medium to remove LB components that could disrupt the coculture mutualism (e.g., amino acids) and the kanamycin to which *R. palustris* was not resistant. Second, all cultures were inoculated using the same volume of washed KEIO_ML9 library, for an *E. coli* starting density of OD_660_≈0.01. Third, *R. palustris* was inoculated at strain-specific cell densities, such that the initial *E. coli*:*R. palustris* ratio approximated the final species ratio known to occur in each coculture type (1:10 and 1:1 for Nx and NxΔAmtB cocultures, respectively) (31). Fourth, all cultures were incubated under identical conditions (i.e., lying flat at 30°C with horizontal shaking and constant illumination) and were harvested in exponential phase after 4.5–5 cell doublings (**Fig. 1C**). We assessed fitness after 4.5–5 cell doublings to enable discrimination between subtle and strong phenotypes (7). Finally, mutant fitness was assessed using 4-fold biological replication for each culture type. All RB-TnSeq experiments in this study met the quality-control standards previously described (**Table S1**) (61).

### Identification of *E. coli* genes affecting fitness in mono- or coculture

The KEIO_ML9 library has a median of 16 unique barcodes (i.e., strains) per gene (61). A fitness score for each *E. coli* strain was calculated as the log_2_ difference in barcode abundance between the washed KEIO_ML9 inoculum (T_0_) and the population after 4–4.5 generations in the mono- or coculture condition. The fitness scores for all strains with insertional disruptions of a given gene were then used collectively to calculate a fitness score for that gene. Genes with no impact on fitness in a given condition had fitness scores close to zero. Negative fitness scores reflected genes that were beneficial to fitness (i.e., gene disruption resulted in a fitness disadvantage), and positive fitness scores indicated genes that were detrimental to fitness (i.e., gene disruption resulted in a fitness advantage). A moderated *t* statistic (*t*-score) was used to estimate how reliably a fitness value differed from zero (61). For each gene, we averaged the fitness values and *t*-scores across culture replicates and defined a strong fitness effect as |mean fitness|>1 and |mean *t*-score|>3.

Of the 3,728 protein-coding genes covered in the KEIO_ML9 library, we could calculate fitness scores for 3,564 genes (95.6% of covered genes) (**Table S2**). Most genes had a negligible impact on fitness (**Fig. S1**). A total of 306 unique genes strongly affected fitness either negatively or positively in one or multiple culture types (**Table S3**). The largest number of fitness-affecting genes was identified in cocultures with *R. palustris* Nx, with 167 genes having negative fitness scores and 105 having positive fitness scores (**Fig. 2A**). Fewer genes affected *E. coli* fitness in coculture with the 3X-NH_4_^+^-excreting *R. palustris* NxΔAmtB, with 122 gene insertions negatively affecting fitness and 2 insertions positively affecting fitness. Only 117 genes were identified to affect *E. coli* monoculture fitness, all of which had negative scores (**Fig. 2A**). Principal component analysis showed clear clustering according to culture type, with the first two principal components explaining approximately 40% of the variability (**Fig. 2B**), and the *E. coli* fitness profiles were more similar between the two types of cocultures than between the monoculture and either coculture (**Fig. 2C, Fig. S2**). Additionally, fitness effects were generally stronger in Nx coculture than in NxΔAmtB coculture (**Fig. 2C**), as we originally hypothesized given that NH_4_^+^ cross-feeding levels are more favorable for *E. coli* growth in NxΔAmtB coculture than in Nx coculture.

**Fig. 2.**
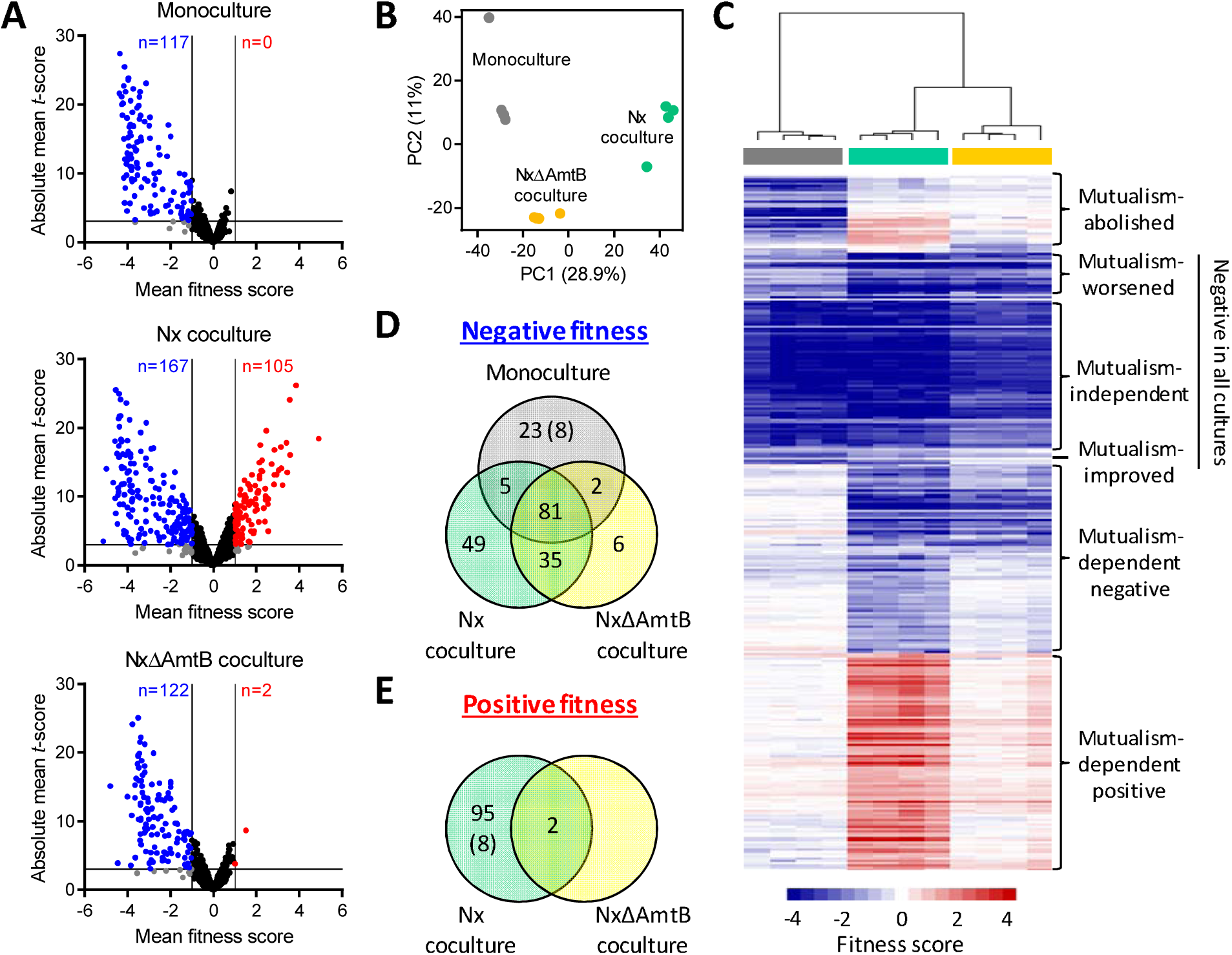
Summary of RB-TnSeq results identifying *E. coli* fitness determinants in monoculture or coculture with *R. palustris*. (A) Volcano plots showing mean fitness scores and absolute mean *t*-scores for 3,564 *E. coli* gene mutants grown in monoculture (top), Nx coculture (middle), or NxΔAmtB coculture (bottom). Each circle represents a single gene, with fitness scores and *t*-scores averaged across culture replicates (n=4). Vertical lines indicate a fitness score threshold of |fitness|>1. Horizontal lines indicate a *t*-score threshold of |*t*-score|>3. Genes with strong negative or positive fitness scores are indicated in blue or red, respectively. Black circles indicate genes that did not meet the fitness score threshold; gray circles indicate genes that met the fitness score threshold but not the *t*-score threshold. (B) Two-component principal component (PC) analysis using fitness scores for 3,564 *E. coli* genes for individual monocultures (gray), Nx cocultures (teal), or NxΔAmtB cocultures (yellow). Percentages indicate variance captured by each of the first two PCs. (C) Heatmap showing fitness scores and designated effect categories for the 306 genes having strong fitness effects in at least one culture type. Genes and samples were clustered hierarchically (average linkage, uncentered correlation). Columns are fitness scores from independent replicates, with culture type indicated by the colored bars below the cluster tree (gray, monoculture; teal, Nx coculture; yellow, NxΔAmtB coculture). (D, E) Venn diagrams of genes having strong negative (D) or positive (E) fitness scores in the indicated culture types. Eight genes had negative fitness scores in monoculture but positive fitness scores in Nx coculture (indicated by parentheses).

Of the 306 genes identified as having strong fitness effects, 81 genes had negative fitness scores in all three cultures (**Fig. 2D**). Based on the magnitudes of effect in the different cultures, these 81 genes fell into three subgroups (**Fig. 2C**). Genes that had similar fitness scores in all three cultures (e.g., *thrA, pfkA*) were considered to have environment-dependent but “mutualism-independent” fitness effects. Genes with fitness scores that were more severe in one or both cocultures than in monoculture (e.g., *lrp, clpP*) were deemed to have “mutualism-worsened” fitness scores. Conversely, genes with negative fitness scores that were less severe in one or both cocultures than in monoculture (e.g., *ilvY, hisI*) were considered to have “mutualism-improved” negative fitness scores (**Fig. 2C**). For simplicity, these 81 genes were classified collectively as “negative in all cultures” for later analyses (**Table S3**).

Separately, 35 genes had negative fitness scores in both cocultures but had negligible fitness effects in monoculture (**Fig. 2D**). An additional 49 genes had negative fitness scores in Nx cocultures only, and an additional six genes had negative fitness scores in NxΔAmtB cocultures only (**Fig. 2D**). Together, we classified these 90 genes as having “mutualism-dependent negative” fitness scores (**Fig. 2C, Table S3**). Notably, 31 genes had negative fitness scores in monoculture but either no fitness effect (23 genes) or a positive fitness score (8 genes) in coculture (**Fig. 2D,E**). These genes were classified as exhibiting “mutualism-abolished fitness defects” (**Fig. 2C, Table S3**). An additional 97 genes had no fitness effect in monoculture but had positive fitness scores in one or both cocultures (**Fig. 2E**) and were thus classified as having “mutualism-dependent positive” fitness scores (**Fig. 2C, Table S3**). Finally, there were seven genes that had negative fitness scores in monoculture and one type of coculture (**Fig. 2C, Table S3**); given their scarcity and their ambiguous fitness effects, we did not designate a label for these seven genes.

As initial validation of our approach, we examined the fitness scores for genes that were previously tested in coculture. Six genes that have no impact on *E. coli* fitness in coculture (*ddpA, ddpX, rutA, argT, patA*, and *potF*) (37) were also shown by RB-TnSeq to have negligible effects in coculture (**Table S2**). Moreover, *amtB* and *ntrC*, which were are crucial for *E. coli* fitness in coculture (37, 39), had negligible impact on monoculture fitness but had two of the top three mutualism-dependent negative fitness scores (**Table 1**). Thus, RB-TnSeq could accurately distinguish between genes that did and did not affect *E. coli* fitness in coculture.

**Table 1.**
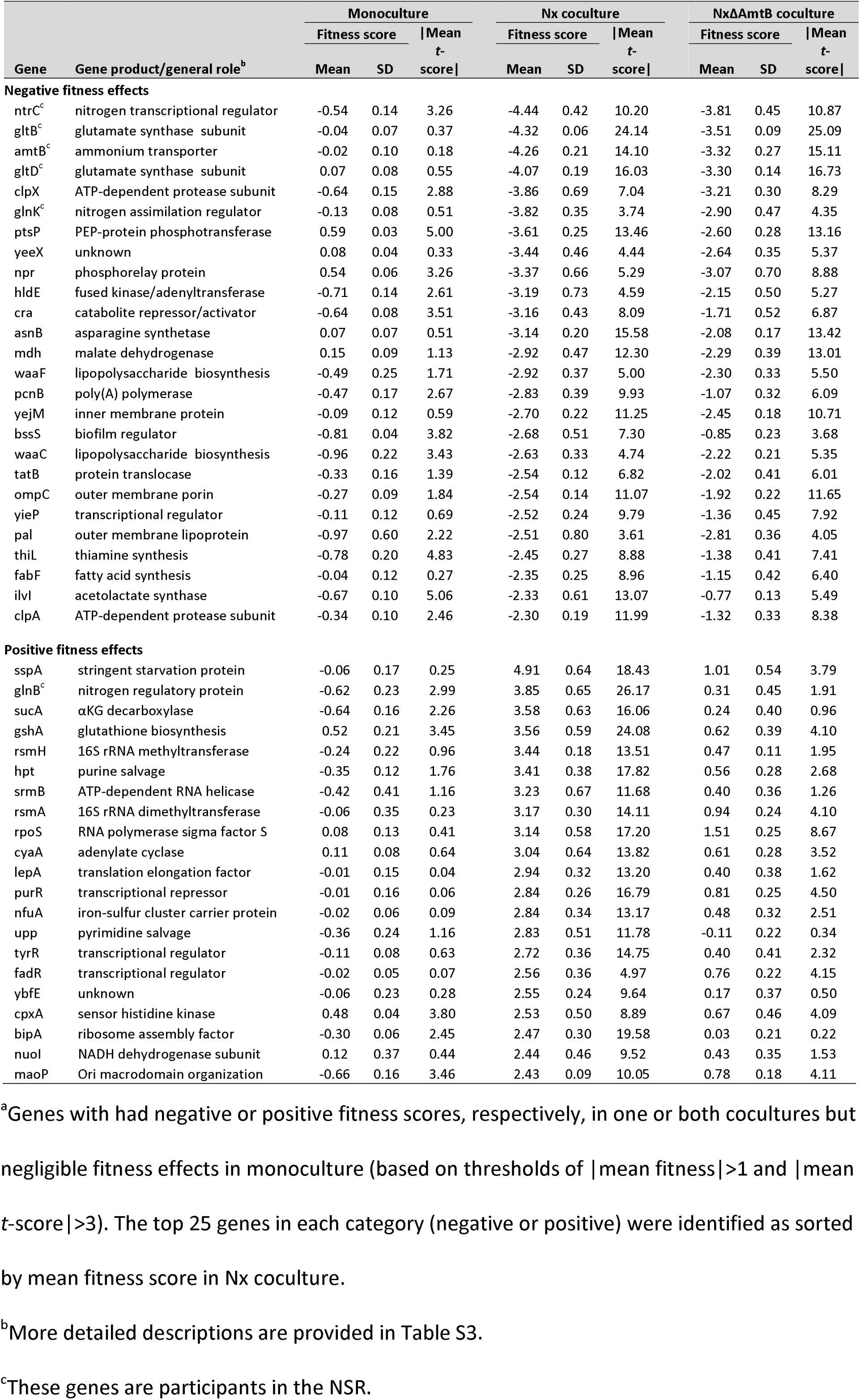
Top 50 genes with mutualism-dependent negative or positive fitness scores^a^.

### Functional classification of *E. coli* genes affecting fitness in mono- or coculture

To investigate the cellular activities that influenced *E. coli* fitness in mono- and coculture, we functionally classified the genes with strong fitness effects using Clusters of Orthologous Groups (COGs) (14). Overall, the 306 genes represented diverse functions, although almost half (48.7%) were involved in metabolism (**Fig. 2A**). Most of the 306 genes pertained to 16 functional categories, with “Amino acid metabolism and transport” predominating, and 67 genes had multiple or poorly defined functions (**Fig. 2B**). Six COG categories were not represented in our screen: “RNA processing and modification”, “Chromatin structure and dynamics”, “Cell motility”, “Nuclear structure”, “Defense mechanisms”, and “Cytoskeleton”. When we examined functional classes according to the fitness effect categories described above, there were clear differences in predominant functions among the different groups. Of the 81 genes with negative fitness scores in all cultures, 55 (67.9%) were related to metabolism (**Fig. 3C**), including 42 of the 65 “Amino acid metabolism and transport” genes in the full 306-gene set (**Fig. 3B,D**). This was unsurprising considering that many amino acid synthesis genes are expected to be essential in the minimal medium we used, which lacks amino acids. Metabolism-related genes accounted for 31.1% and 38.1% of the mutualism-dependent negative and positive groups, respectively (**Fig**. 3C), but these similar proportions were due to distinct gene classes. In the mutualism-dependent negative group, 19 (67.8%) of the 28 metabolism-related genes were involved in “Amino acid metabolism and transport” (n=13) and “Inorganic ion transport and metabolism” (n=6) (**Fig. 3D**). In contrast, the metabolism-related genes in the mutualism-dependent positive group were predominantly linked to “Energy production and conversion” (16 of 37 genes, 43.2%), with the remaining genes being fairly evenly spread among other metabolic functions (**Fig. 3D**). The prevalence of genes related to other general functions was also distinct between the mutualism-dependent negative and positive groups (**Fig. 3C**). The mutualism-dependent negative group had the largest proportion of genes related to “Cellular processes and signaling” among the groups (27.8%) (**Fig. 3C**), and these were primarily involved in “Cell wall and membrane biogenesis” (n=13), “Transcription” (n=8), and “Signal transduction” (n=6) (**Fig. 3D**). This group also had the largest proportion of “Poorly characterized” genes among the groups (14.4%) (**Fig. 3C**). Meanwhile, the mutualism-dependent positive group had a smaller proportion of genes involved in “Cellular processes and signaling” (19.6%) and had the largest proportion of genes related to “Information storage and processing” (17.5%) among the groups (**Fig. 3C**). Finally, the mutualism-abolished fitness defect group consisted almost entirely (83.9%) of metabolism-related genes (**Fig. 3C**). The largest proportion of these genes was involved in “Nucleotide metabolism and transport” (25.8%), followed by “Coenzyme metabolism” and “Carbohydrate metabolism” (16.1% each) (**Fig. 3D**); “Nucleotide metabolism and transport” and “Coenzyme metabolism” were poorly represented (1.2–7.4%) in the other groups. Overall, these data indicate that genes involved in diverse cellular functions impact *E. coli* fitness in coculture with *R. palustris*.

**Fig. 3.**
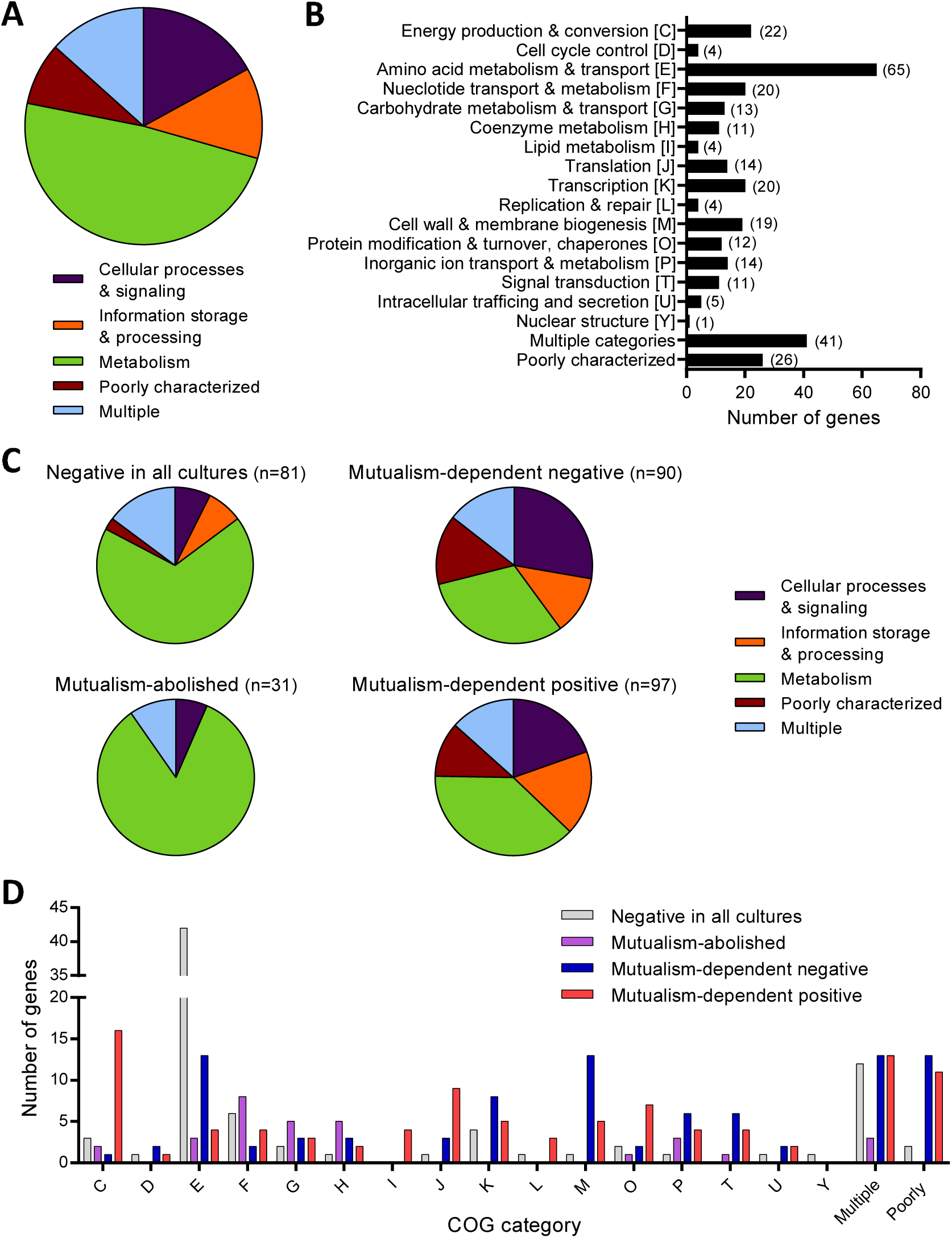
Functional summary of genes exhibiting strong fitness effects in one or more culture type. (A-D) Functional categories, based on COGs, for (A,B) all 306 genes with |mean fitness score|>1 and |mean *t*-score|>3 and (C, D) gene subsets exhibiting the designated fitness effects. Genes listed as “Poorly characterized” include genes with general functional predictions only (COG category R), genes with unknown function (COG category S), and genes with no COG classification. Genes listed as “Multiple” have more than one COG category assignment. Gene names, descriptions, fitness scores, *t*-scores, and COG classifications are provided in Table S3. COG categories are the same in panels B and D. (B) Numbers in parentheses denote the number of genes in each category. (A, C) The pie charts in panels A and C represent the same genes in B and D, respectively, but with COG categories grouped into generalized cell functions, as follows: Cellular processing and signaling, COG categories DMNOTUVYZ; Information storage and processing, COG categories ABJKL; and Metabolism, COG categories CEFGHIPQ.

### Only a portion of the NSR benefits *E. coli* fitness in coculture

As mentioned above, the transcriptional regulator NtrC and the NH_4_^+^ transporter AmtB were previously shown to be decisive *E. coli* fitness determinants in coculture (37, 39), and our TnSeq data agreed with these results (**Table 1**). NtrC and AmtB are central players in the NSR (57). Other NSR-related genes also exhibited mutualism-dependent fitness effects in our screen (**Fig. 4A**). In fact, NSR-related genes comprised five of the top seven mutualism-dependent fitness scores (**Table 1**). These data further authenticated the importance of the *E. coli* NSR for mutualistic coexistence with *R. palustris*. However, not all NSR genes contributed to *E. coli* fitness in coculture. For example, the nitrogen assimilation control (*nac*) gene had limited impact on fitness in any culture type (mean fitness, 0.20 for monoculture, −0.58 for Nx coculture, −0.48 for NxΔAmtB coculture) (**Fig. 4A**). Nac is the second main transcriptional regulator of the NSR, behind NtrC, and of the 23 operons within the NtrC regulon, which includes nac, nine are regulated indirectly via Nac (66). *E. coli nac* expression is upregulated >40-fold in coculture compared to monoculture (37). However, only four Nac-regulated genes are differentially expressed in coculture versus monoculture (37), and none of these met our thresholds for strongly impacting fitness in coculture (**Table S4**). Thus, although the low NH_4_^+^ cross-feeding level in coculture activates the NSR and the resulting NtrC-mediated induction of some NSR genes is crucial for *E. coli* fitness, the Nac-mediated arm of the *E. coli* NSR appears to be dispensable for mutualistic coexistence with *R. palustris*.

**Fig. 4.**
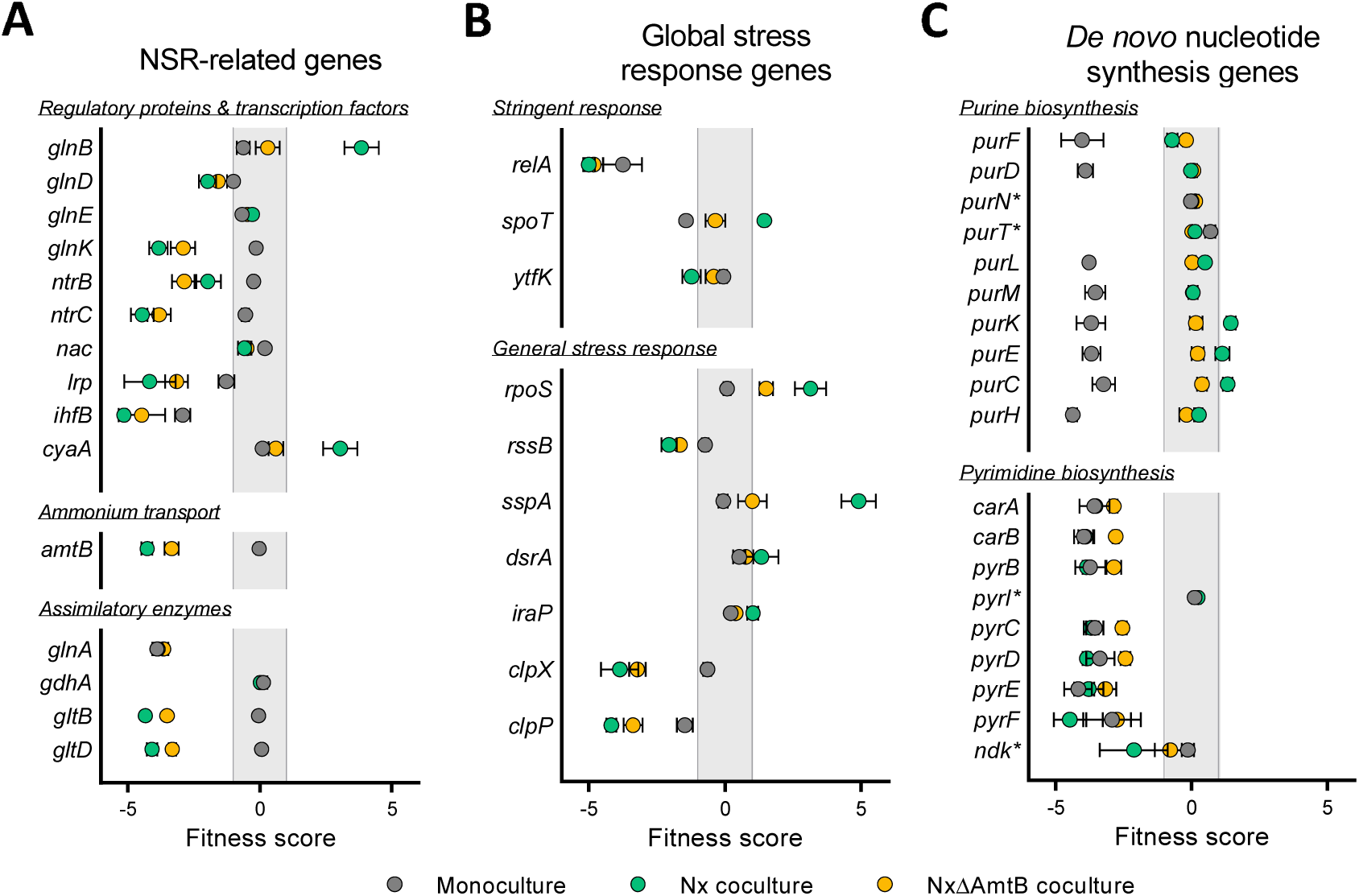
Fitness scores for select *E. coli* genes. (A-C) Mean fitness scores of select genes involved in (A) the NSR, (B) global stress responses, and (C) *de novo* nucleotide biosynthesis for *E. coli* grown in monoculture (gray), Nx coculture (teal), or NxΔAmtB coculture (yellow). Error bars indicate standard deviations (n=4). (C) Asterisks denote genes that are known to have redundant functions or be dispensable for *de novo* synthesis.

Notably, whereas disruption of many NSR genes negatively affected fitness in coculture, *glnB* had a strong positive fitness score in Nx coculture (**Table 1, Fig. 4A**). In response to nitrogen availability, GlnB (also known as PII) regulates glutamine synthetase (GS) adenylation and NtrC phosphorylation (44). In the absence of GlnB, the cell cannot sense when there is ample nitrogen and the NSR is activated via NtrC even under high-nitrogen conditions (57). We posit that the *glnB* mutant exhibits better fitness upon introduction into coculture due to constitutively active NtrC. Specifically, when the KEIO_ML9 library was initially grown in LB, NtrC (and the NSR) would be inactive in most cells due to abundant nitrogen. However, in the *glnB* mutant, constitutively active NtrC would be activating transcription of the NSR genes, including *amtB* (44). Consequently, unlike the rest of the population, the *glnB* mutant would already have high levels of AmtB upon inoculation in coculture and therefore be primed for mutualistic growth. Although possible, we reason it unlikely that the *glnB* fitness effect is mediated via GS adenylation because disruption of *glnE*, the adenylyltransferase responsible for GS adenylation, had a negligible fitness effect in all culture types (**Fig. 4A**). Future studies will be needed to verify how GlnB affects fitness in coculture.

### Metabolic pathway choice influences *E. coli* fitness in coculture

The second and fourth top mutualism-dependent negative fitness scores went to *gltB* and *gltD* (**Table 1**). These genes are NSR-related but their effect was likely assimilatory rather than regulatory. *gltB* and *gltD* encode the two subunits of glutamate synthetase (GOGAT), which is one of two *E. coli* glutamate synthesis enzymes. GOGAT transfers an amino group from glutamine to α-ketoglutarate (αKG), resulting in two molecules of glutamate (20). In the other pathway, glutamate is synthesized directly from αKG and NH_4_^+^ by glutamate dehydrogenase (GDH), encoded by *gdhA*. The use of GOGAT or GDH depends on NH_4_^+^ concentrations, with GOGAT being important under low-NH_4_^+^ conditions due to a much higher affinity for NH_4_^+^ (20). The *gdhA* gene exhibited negligible fitness effects under any condition (**Fig. 4A**), presumably because GOGAT can compensate for the absence of GDH under both low- and high-NH_4_^+^ conditions (57). In contrast, *gltB* and *gltD* had negative fitness scores in both cocultures (**Fig. 4A**), with the score magnitudes inversely correlating with NH_4_^+^ cross-feeding levels. These data suggest that the low NH_4_^+^ levels in coculture compel *E. coli* to use GOGAT for mutualistic growth.

To directly assess the effect of GOGAT on growth in coculture, we grew the KEIO Δ*gltB* mutant in both monoculture and Nx coculture. We used the KEIO Δ*fimA* mutant as a control strain, as *fimA* had a negligible impact on fitness (|mean fitness|<0.1) in all culture types (**Table S2**). As expected, the Δ*fimA* mutant exhibited robust growth in both monoculture and Nx coculture (**Fig. 5A**), with comparable growth trends to those observed with strain MG1655 (31). In agreement with the RB-TnSeq data, the Δ*gltB* mutant grew well in monoculture, matching the Δ*fimA* mutant growth trends (**Fig. 5A**). In contrast, Δ*gltB* Nx cocultures exhibited an extended lag period of ∼85 h, versus ∼26 h in Δ*fimA* Nx cocultures (**Fig. 5B**). The *gltB* RB-TnSeq fitness score near −5 in Nx cocultures, which indicates negligible growth, came from samples harvested at ∼54 h, which would be within this extended lag period. From these results, we conclude that GOGAT is important for *E. coli* fitness in coculture.

**Fig. 5.**
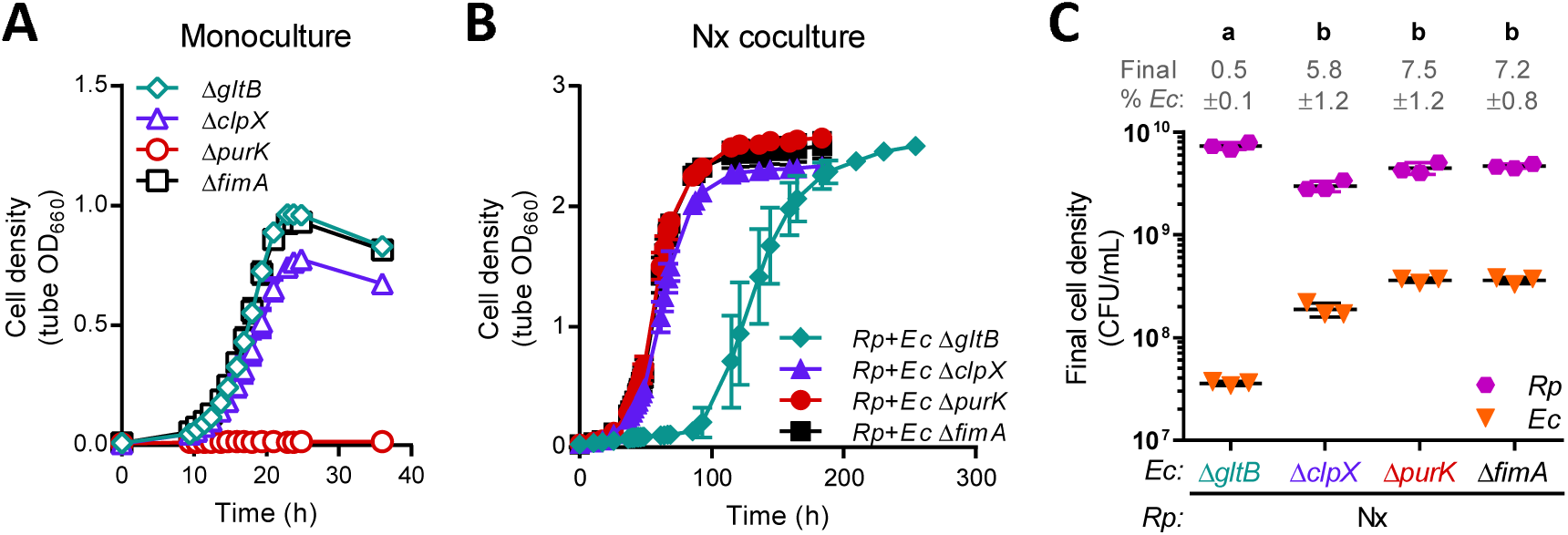
Growth of select KEIO mutants in monoculture and Nx coculture. (A,B) Growth curves of individual KEIO mutants in monoculture (A) or coculture with *R. palustris* Nx (B). (C) Final cell densities of individual *E. coli* KEIO mutants and *R. palustris* Nx after growth in coculture. CFUs were measured at the final time points in the respective growth curves in panel B. Different letters indicate statistical differences between groups (P<0.05, one-way analysis of variance with Tukey’s posttest, n=3).

Nonetheless, we were intrigued by the eventual growth of Δ*gltB* Nx cocultures. The abrupt growth after an extended lag phase (**Fig. 4C**) suggested that the absence of GOGAT was overcome in some manner. When we examined final cell densities, we found that the proportion of the Δ*gltB* mutant within Nx cocultures was >10-fold less than that of the Δ*fimA* mutant, resulting from both lower *E. coli* cell densities and higher *R. palustris* cell densities (**Fig. 4C**). Thus, one explanation is that the skewed species ratio in Δ*gltB* Nx coculture provided higher NH_4_^+^ levels per *E. coli* cell, thereby allowing GDH-mediated glutamate synthesis and coculture growth; the extended lag could represent the period during which the coculture was reaching this alternate species ratio. It is also possible that the Δ*gltB* mutant acquired suppressor mutations that conferred elevated GDH activity, which can rescue GOGAT mutant growth under low NH_4_^+^ conditions (63). Overall, enzyme choice for glutamate synthesis clearly affects *E. coli* fitness during mutualistic growth, as GDH does not compensate for the absence of GOGAT in coculture.

### Multiple global regulatory systems influence *E. coli* fitness in coculture

The NSR is the primary cellular response to nitrogen limitation in *E. coli*. However, the *E. coli* NSR is coupled to the stringent response under low-nitrogen conditions because NtrC directly activates transcription of *relA* (6). *E. coli* RelA is the primary enzyme for the synthesis of the alarmone (p)ppGpp; the remaining (p)ppGpp is synthesized by the (p)ppGpp synthase/hydrolase SpoT (19). Accumulation of (p)ppGpp coordinates the redistribution of resources to enable continued growth and survival amid nutrient stress (19). Both *relA* and *spoT* were identified by RB-TnSeq as strongly influencing *E. coli* fitness in one or multiple culture types (**Fig. 4B**). *relA* had negative fitness scores in all cultures, but the effects were stronger in cocultures than in monoculture. In contrast, *spoT* had a negative fitness score in monoculture but a positive fitness score in Nx coculture and a negligible effect in NxΔAmtB coculture. SpoT is essential in *relA*^+^ *E. coli* because uncontrolled, high levels of (p)ppGpp inhibit growth (54). However, viable *spoT* insertion mutants have been identified by both RB-TnSeq (50) and transposon-directed insertion site sequencing (16). It is possible that the RB-TnSeq *spoT* strains harbor compensatory secondary mutations that enable growth (42, 56), and thus we mention the *spoT* data for transparency only. Given that (i) *E. coli* (p)ppGpp levels increase in response to nitrogen limitation (59), (ii) RelA mutants are severely impaired for the stringent response (12), and (iii) disruption of *relA* was associated with a negative fitness effect that was exacerbated in coculture (**Fig. 4B**), we infer that the *E. coli* stringent response is activated in coculture and contributes to fitness during mutualistic growth.

As (p)ppGpp affects the expression of ∼500 *E. coli* genes (12), we can currently only speculate as to which stringent response targets might affect fitness in coculture. Several (p)ppGpp-regulated genes differentially affected *E. coli* fitness in coculture versus monoculture, including *upp, nuoA–N, cyoDE, sucACD, ptsG*, and *plsB* (**Table S3**). Most intriguing to us, three interlinked global regulators that are influenced by (p)ppGpp were also hit in our screen: leucine responsive protein (Lrp), integration host factor (IHF), and the alternative sigma factor RpoS (**Fig. 4A,B**). Lrp affects transcription of ∼236 *E. coli* genes and coordinates multiple regulatory networks to adjust cellular metabolism in response to environmental perturbations (8). Lrp is also considered part of the nitrogen assimilation network because its regulon includes *gltBD* (57). The second regulator, IHF, has many functions including serving as an important NSR transcription factor (57) and regulating anaerobic fermentative metabolism in *E. coli* (26). The expression of both Lrp and IHF are upregulated by the stringent response (3, 12), and disruption of *lrp* or *ihfB* resulted in mutualism-worsened fitness (**Fig. 4A**). Lrp and IHF might be beneficial in coculture by activating enzymes like GOGAT, but their effects could also be unrelated to the NSR given the diverse genes in their regulons. The third global regulator, RpoS (σ^38^), mediates the general stress response and directly or indirectly regulates ∼10% of all *E. coli* genes (5). The general stress response is stimulated by nitrogen starvation (17), although not via NtrC (35), and like Lrp and IHF, RpoS levels increase with (p)ppGpp levels (15). Unlike disruption of *lrp* and *ihfB*, however, disruption of *rpoS* resulted in enhanced fitness in both cocultures (**Fig. 4B**). Thus, the general stress response appears to be detrimental to *E. coli* fitness in coculture.

Lending support to this notion, several regulators of RpoS exhibited negative or positive mutualism-dependent fitness effects in correlation with the gene’s expected effect on RpoS levels (**Fig. 4B**). For example, the protease ClpXP, in conjunction with the adapter protein RssB, degrades RpoS (5). Disruption of any of these genes results in RpoS accumulation (49, 64), and all three had negative fitness scores in coculture (**Fig. 4B**). Because *clpX* had the fifth most negative mutualism-dependent fitness score (**Table 1**), we examined KEIO Δ*clpX* mutant growth in both monoculture and Nx coculture. Whereas RB-TnSeq indicated *clpX* to have a minor fitness effect in monoculture but a major effect in coculture (**Fig. 4B**), the KEIO Δ*clpX* mutant exhibited comparable minor growth defects in both culture types, as evidenced by lower final cell densities, lower growth yields, and slightly lower growth rates compared to control Δ*fimA* cultures (**Fig. 5A,B, Fig. S3**). We hypothesize that the exacerbated negative fitness effect of *clpX* in RB-TnSeq cocultures is due to competition against the other *E. coli* mutants in the population. In other words, engaging in a mutualistic interaction with a partner species could amplify or dampen gene-specific fitness effects stemming from intraspecific competition.

Regardless of potential intraspecific competition effects on *clpX* mutant growth trends, the general stress response could hamper *E. coli* fitness in coculture by several mechanisms. First, RpoS could directly control expression of target genes that influence *E. coli* fitness. Alternatively, because sigma factors compete for a limited amount of RNA polymerase (45), induction of the general stress response might limit gene expression driven by other sigma factors important in coculture, such as RpoN. RpoN drives expression of many NSR genes, including *ntrC* (57), and has also been implicated in the control of membrane and cell wall biogenesis (13), a function that was well represented by genes exhibiting mutualism-dependent fitness effects (**Fig. 3D**). Notably, *E. coli* sigma factor competition is influenced by ppGpp (24), Lrp (60), and IHF (10), as well as two other genes hit in our screen, namely *ptsN* (32) (**Table 2**), encoding part of the regulatory PTS^Ntr^ system (47), and *cyaA* (10) (**Table 1**), encoding the cAMP-synthesizing adenylate cyclase. Thus, sigma factor competition resulting from the activation of multiple stress responses might curtail *E. coli* fitness in coculture.

**Table 2.** Top 25 genes exhibiting mutualism-abolished fitness defects^a^.

| Gene | Gene product/general role<sup>b</sup> | Monoculture |  |  | Nx coculture |  |  | NxΔAmtB coculture |  |  |
| --- | --- | --- | --- | --- | --- | --- | --- | --- | --- | --- |
|  |  | Fitness score |  | \|Mean *t*-score\| | Fitness score |  | \|Mean *t*-score\| | Fitness score |  | \|Mean *t*-score\| |
|  |  | Mean | SD |  | Mean | SD |  | Mean | SD |  |
| purH | purine biosynthesis | -4.38 | 0.17 | 21.62 | 0.27 | 0.15 | 2.14 | -0.18 | 0.27 | 1.53 |
| sucD | succinyl-CoA synthetase subunit | -4.14 | 0.76 | 5.69 | -0.93 | 0.26 | 2.45 | -0.44 | 0.13 | 1.24 |
| purF | purine biosynthesis | -4.02 | 0.78 | 11.85 | -0.71 | 0.19 | 3.35 | -0.20 | 0.11 | 1.18 |
| purD | purine biosynthesis | -3.91 | 0.28 | 16.73 | -0.02 | 0.13 | 0.06 | 0.08 | 0.13 | 0.55 |
| sucC | succinyl-CoA synthetase subunit | -3.81 | 0.11 | 10.47 | -0.89 | 0.43 | 4.51 | -0.14 | 0.15 | 0.79 |
| purL | purine biosynthesis | -3.77 | 0.06 | 18.86 | 0.50 | 0.09 | 3.70 | 0.03 | 0.13 | 0.21 |
| purK | purine biosynthesis | -3.70 | 0.53 | 13.84 | 1.45 | 0.17 | 9.73 | 0.17 | 0.24 | 1.13 |
| purE | purine biosynthesis | -3.68 | 0.34 | 6.70 | 1.13 | 0.26 | 5.00 | 0.23 | 0.22 | 0.96 |
| purM | purine biosynthesis | -3.54 | 0.38 | 9.72 | 0.06 | 0.16 | 0.31 | 0.05 | 0.07 | 0.21 |
| purC | purine biosynthesis | -3.23 | 0.42 | 4.95 | 1.32 | 0.18 | 4.24 | 0.39 | 0.19 | 1.44 |
| pdxJ | vitamin B6 biosynthesis | -3.06 | 0.44 | 8.89 | -0.91 | 0.26 | 2.83 | -0.52 | 0.11 | 2.19 |
| pdxA | vitamin B6 biosynthesis | -2.95 | 0.09 | 15.11 | -0.94 | 0.40 | 5.73 | -0.18 | 0.15 | 1.36 |
| eda | KHG/KDPG aldolase | -2.78 | 0.33 | 4.25 | 1.32 | 0.35 | 3.88 | -0.51 | 0.38 | 1.22 |
| gcvR | transcriptional regulator | -2.76 | 0.35 | 8.30 | -0.63 | 0.16 | 2.92 | -0.53 | 0.22 | 2.63 |
| ptsG | glucose transport | -2.55 | 0.26 | 13.72 | 1.00 | 0.23 | 7.38 | 0.11 | 0.08 | 0.78 |
| pdxB | vitamin B6 biosynthesis | -2.51 | 0.29 | 12.70 | -0.71 | 0.42 | 4.03 | -0.09 | 0.03 | 0.63 |
| argD | arginine/ornithine biosynthesis | -1.98 | 0.11 | 15.35 | -0.50 | 0.04 | 4.07 | -0.25 | 0.05 | 2.14 |
| ptsN | phosphotransferase system enzyme | -1.84 | 0.12 | 7.28 | 0.91 | 0.32 | 4.59 | -0.74 | 0.18 | 3.33 |
| pgl | 6-phosphogluconolactonase | -1.49 | 0.61 | 4.46 | 1.38 | 0.21 | 5.37 | 0.04 | 0.26 | 0.17 |
| spoT | (p)ppGpp synthase/hydrolase | -1.43 | 0.07 | 6.83 | 1.45 | 0.12 | 8.32 | -0.35 | 0.36 | 1.75 |
| nadA | NAD biosynthesis | -1.39 | 0.24 | 6.90 | -0.07 | 0.17 | 0.41 | -0.03 | 0.07 | 0.15 |

<sup>a</sup>Genes had negative fitness scores in monoculture but negligible effects or positive fitness scores in one or both cocultures (based on our thresholds of |mean fitness|>1 and |mean *t*-score|>3). The top 25 genes were identified as sorted by mean fitness score in monoculture.
<sup>b</sup>More detailed descriptions are provided in Table S3.

### *E. coli* mutants defective for *de novo* purine biosynthesis are rescued in coculture with *R. palustris*

Among the hits from our screen, we were most intrigued by the genes exhibiting mutualism-abolished fitness defects (**Table 2**) because, unlike some other coculture systems (18, 46, 65), our coculture does not enforce coexistence through an engineered auxotrophy. One mechanism by which coculturing could restore *E. coli* mutant fitness is by slowing the growth of all *E. coli* strains. In our coculture, the *E. coli* growth rate is dictated by the NH_4_^+^ cross-feeding level (31). Thus, *E. coli* mutants with growth rate defects in monoculture could have higher fitness scores in coculture simply due the nitrogen-limiting conditions that slow the growth of all *E. coli* mutants in the competition. Indeed, one of the genes exhibiting mutualism-abolished fitness defects that could be explained by slower growth of competing strains was *ptsG* (**Table 2**), which encodes the permease of the glucose phosphotransferase transport system. Disruption of *ptsG* in *E. coli* results in poor glucose uptake and slower growth on glucose minimal medium (48). This mutation would have little consequence in coculture as long as the decreased growth rate of the *ptsG* mutant was equivalent or faster than the growth rate dictated by the NH_4_^+^ level. Notably, the positive fitness score of *ptsG* in Nx coculture (**Table 2**) indicates that the mutant grew better than the average *E. coli* strain in the population, perhaps due to directing carbon more efficiently to biomass (48). An alternative mechanism by which coculturing could restore fitness of an *E. coli* mutant is by decreasing the importance of a given metabolic pathway. For example, nitrogen starvation results in decreased glucose uptake rates in *E. coli* (9), potentially further reducing the importance of *ptsG* in coculture.

However, most of the top 25 genes exhibiting mutualism-abolished fitness defects were not readily explained by slowed growth of all *E. coli* strains, as the genes were involved in synthesizing essential cellular building blocks, such as purines (**Table 2**). Of the 14 *E. coli* genes involved in AMP and GMP synthesis, *purA, purB, guaA*, and *guaB* were not in our screen. Eight of the 10 remaining genes had severe negative fitness scores in monoculture but had negligible effects or positive fitness scores in coculture (**Fig. 3C**). The two genes that did not exhibit fitness defects in monoculture, *purN* and *purT*, are redundant in function (36) and thus would not have a phenotype when mutated individually. Importantly, *de novo* purine synthesis mutants are purine auxotrophs (25). Thus, the success of the *pur* mutants specifically in coculture suggested that coculturing alleviated purine auxotrophy. Coculturing did not eliminate the fitness defects of *de novo* pyrimidine synthesis mutants (**Fig. 4C**), indicating that coculture-mediated rescue of nucleotide auxotrophy was specific to purines.

To verify that coculture with *R. palustris* could rescue *E. coli* purine auxotroph growth, we tested growth of the KEIO Δ*purK* mutant in monoculture and Nx coculture. As expected, the Δ*purK* mutant did not grow in monoculture (**Fig. 5A**), as no purines were provided. However, in agreement with the RB-TnSeq data, Δ*purK* Nx cocultures exhibited comparable growth trends to those of control Δ*fimA* Nx cocultures (**Fig. 5B,C** and **Fig. S3**). These data demonstrate that coculture with *R. palustris* can restore *E. coli* purine auxotroph growth to wild-type levels. In support of this conclusion, *E. coli pur* gene transcription is downregulated in coculture with *R. palustris* (37). Although this downregulation was originally ascribed to slower *E. coli* growth (37), *de novo* purine synthesis is repressed by the presence of purines in the medium (29), as could occur if provided by *R. palustris*.

## CONCLUSION

Using RB-TnSeq, we identified *E. coli* genes that were beneficial or detrimental to *E. coli* fitness in coculture with *R. palustris*. This rich dataset provided both general gene fitness trends and some specific molecular insights into the mutualistic relationship, such as the importance or lack thereof of various regulatory and metabolic pathways and the existence of unexpected purine cross-feeding. Similar to findings from TnSeq experiments on cocultures mimicking coinfections (2, 23, 33), we observed that diverse cellular functions impact *E. coli* fitness in coculture with *R. palustris*. These observations underscore that the genetic and physiological architectures underlying microbial interactions, cooperative or otherwise, are complex and often difficult to predict, even in engineered systems. However, architectures common to microbial interactions may become apparent as additional genome-scale fitness studies of microbial consortia are performed. For example, cell wall and membrane biogenesis impacted coli fitness in coculture with *R. palustris*, and membrane transport was also important for *E. coli* growth in a cheese rind community (43), suggesting that membrane-related functions may be of broad importance for *E. coli* growth in the presence of additional species. Our work further demonstrates that experimentation on microbial consortia will invariably unmask hidden, but nonetheless important, aspects of microbial physiology.

## MATERIALS AND METHODS

### Strains and media

All *R. palustris* strains were derived from the wild-type strain CGA009 (30). *R. palustris* Nx (CGA4005) harbors (i) a nifA* allele, resulting in constitutive nitrogenase expression and NH_4_^+^ excretion, (ii) Δ*hupS*, which prevents H_2_ uptake, and (iii) Δ*uppE*, which prevents cell-cell aggregation (31). *R. palustris* strain NxΔAmtB (CGA4021) harbors the same three mutations as strain Nx and additional deletions of *amtB1* and *amtB2* (31). All *E. coli* strains were derived from the *E. coli* K12 strain BW25113, the parent strain of the KEIO single gene deletion collection (4). The RB-TnSeq *E. coli* library, KEIO_ML9, was made and characterized previously (61). The single-gene deletion KEIO mutants used in this study were: JW4277 (Δ*fimA*::Km), JW379 (Δ*gltB*::Km), JW0428 (Δ*clpX*::Km), and JW0511 (Δ*purK*::Km) (4). KEIO mutant genotypes were confirmed by PCR.

For routine cultivation, *R. palustris* colonies were isolated on defined mineral (PM) (28) agar supplemented with 10 mM succinate and 0.1% (w/v) yeast extract (PMsuccYE). Plates were incubated anaerobically in a jar with a GasPack sachet (BD, Franklin Lakes, NJ, USA) at 30°C. *E. coli* KEIO mutant colonies were isolated on LB agar containing 30 µg/µL kanamycin and incubated aerobically at 30°C in the dark. All anaerobic plates and monocultures, and all cococultures were illuminated using a 60 W soft white halogen bulb (750 lumens). All anaerobic liquid mono- and cocultures were laid flat and shaken at 150 rpm. Other aspects of the growth conditions for the KEIO_ML9 library are described below.

Anaerobic monocultures and cocultures were cultivated in 10 mL defined coculture medium (MDC) (31) in 27-mL anaerobic test tubes flushed with 100% N_2_ and sealed with rubber stoppers and aluminum crimps prior to autoclaving. After autoclaving, MDC was supplemented with 25 mM glucose and cation solution (1% [v/v]; 100 mM MgSO_4_ and 10 mM CaCl_2_). Additionally, monocultures received 15 mM NH_4_Cl whereas cocultures received 15 mM NaCl.

### Analytical procedures

Culture growth was assessed by optical density at 660 nm (OD_660_) using a Genesys 20 visible spectrophotometer (Thermo-Fisher). OD measurements for all anaerobic monocultures and cocultures were taken in culture tubes without sampling. Growth rates were calculated using OD values between 0.1–1.0 where cell density and OD_660_ are linearly correlated. Cell densities of aerobic *E. coli* monocultures were measured in cuvettes. All final culture densities were also measured in cuvettes, and samples were diluted as necessary to achieve values <1 OD_660_. High-performance liquid chromatography (Shimadzu) was used to quantify glucose as described (40).

### RB-TnSeq culture preparation, incubation, and harvesting

A 2-mL aliquot of the KEIO_ML9 library, stored in 25% (v/v) glycerol at −80°C, was thawed on ice and used to inoculate 48 mL LB with 30 µg/µL kanamycin in a 250 mL aerobic flask. The culture was incubated at 30°C with shaking at 150 rpm until mid-exponential phase (OD_660_=0.44). Cells from 35 mL of culture were pelleted (4000 rpm, 5 min, room temperature), washed four times with 33 mL MDC, and resuspended in MDC to OD_660_=1.0.

Single colonies of *R. palustris* Nx or NxΔAmtB were inoculated to carbon-limited MDC supplemented with 3 mM acetate. Once the OD of these starter cultures plateaued, the cultures were diluted or concentrated as necessary in MDC to achieve desired cell densities (see below).

The following protocol was used to inoculate three types of cultures, each in quadruplicate: (i) KEIO_ML9 monoculture; (ii) Nx coculture, pairing KEIO_ML9 with *R. palustris* Nx; and (iii) NxΔAmtB coculture, pairing KEIO_ML9 with *R. palustris* NxΔAmtB. Each monoculture and coculture was inoculated with 100 uL washed KEIO_ML9 library. Separately, 1 mL of washed KEIO_ML9 library was centrifuged and the cell pellet (input sample, T_0_) was stored at −80°C for barcode sequencing (BarSeq) analysis. For *R. palustris*, cells from duplicate Nx and NxΔAmtB starter cultures were resuspended to OD_660_=0.6. For *R. palustris* NxΔAmtB, 100 uL of the resuspended cells were used directly to inoculate each NxΔAmtB coculture. For *R. palustris* Nx, 1 mL aliquots of of the resuspended cells were concentrated 10-fold into 100 µL MDC and then inoculated to start each Nx coculture. Each *R. palustris* starter culture replicate was used to inoculate two corresponding cocultures, for a total of four cocultures of each type.

Each culture was allowed to double 4.5-5 times, at which point samples were harvested (1 mL for Nx cocultures; 2 mL for monocultures and NxΔAmtB cocultures) and centrifuged, and the cell pellets were stored at −80°C for BarSeq analysis.

### BarSeq sample preparation and sequencing

Total genomic DNA (gDNA) was isolated from the T_0_, monoculture, and coculture cell pellets using a Wizard Genomic DNA Purification kit (Promega) or Bactozol kit (Molecular Research Center). Because *E. coli* abundance differed between T_0_ and monoculture samples, Nx cocultures, and NxΔAmtB cocultures (100%, ∼10%, and ∼50%, respectively) (31), different amounts of total purified gDNA were used as template in 50 µL BarSeq PCRs, as follows: 100 ng gDNA from T_0_ and monocultures; 300 ng gDNA from Nx cocultures; and 200 ng gDNA from NxΔAmtB cocultures. BarSeq PCRs were performed using the previously described BarSeq98 protocol and primers, which add a unique experimental index to each sample to enable multiplex sequencing (61).

PCR product analysis, pooling, and Illumina sequencing was performed at the Indiana University Center for Genomics and Bioinformatics. The quality and concentration of the amplified PCR products were determined using an Agilent 2200 TapeStation. Equimolar concentrations of PCR product from all samples were pooled, and the pooled PCR products were purified using Agencourt AMpure beads (Beckman Coulter) and then sequenced using a single Illumina NextSeq 75 cycle high-output run.

### Fitness data analysis, COG assignment, and statistical analyses

Gene fitness values were determined from the BarSeq data using the described pipeline (61). Gene fitness scores and *t*-scores were averaged across quadruplicate replicates for each culture type, and genes with |mean fitness score|>1 and |mean *t*-score|>3 in at least one culture type were considered to have strong fitness effects.

COG assignments for the 306 genes exhibiting strong fitness effects were curated manually by sequentially converting gene locus tags to K numbers and then to COG numbers using the files at https://www.genome.jp/dbget-bin/get_linkdb?-t+genes+gn:T00007 and https://www.genome.jp/kegg/files/ko2cog.xl. As this method did not identify COG numbers for all genes, some COG numbers were assigned directly from the locus tags using the JGI Gene Search database (https://img.jgi.doe.gov/cgi-bin/m/main.cgi?section=GeneSearch&page=searchForm). The COG numbers were then used to assign COG categories based on the files available at the following two sites: ftp://ftp.ncbi.nih.gov/pub/COG/COG2014/static/lists/listEsccol.html and ftp://ftp.ncbi.nih.gov/pub/wolf/COGs/COG0303/cogs.csv. Finally, COG categories were tallied using Microsoft Excel.

Principal component analysis was performed in ClustVis (41) using all calculated fitness scores (3,564 genes). Results were plotted using Prism v6 (GraphPad, San Diego, CA, USA). Hierarchical clustering (average linkage, uncentered coordination) was performed in Cluster 3.0 (11), and the resulting data were visualized using Java TreeView (52).

### Mutant fitness validation using KEIO mutants

Overnight cultures of KEIO mutants received glycerol to final concentration of 25% (v/v), and 250 μL aliquots were stored at −80°C to mimic the conditions used for RB-TnSeq experiments. Individual aliquots were thawed on ice and used to inoculate triplicate 6-mL aerobic starter cultures. At mid-exponential phase (OD_660_=0.40–0.55), cultures were pelleted (4000 rpm, 5 min, room temperature), washed four times with 5 mL MDC, and resuspended in MDC to OD_660_=0.5. Triplicate *R. palustris* Nx starter cultures were prepared under carbon-limiting conditions as described above. The Nx cultures were resuspended to OD_660_=0.3, and 1-mL cell aliquots were pelleted (4000 rpm, 5 min, room temperature) for use below.

KEIO mutant monocultures were inoculated using 100 μL washed cells. For cocultures, one cell pellet of each Nx replicate was resupsended in 100 μL of a washed KEIO mutant cell suspension and each resulting 100 μL pairing was used to inoculate one of three Nx coculture replicates. Cell densities (CFU/mL culture) were determined by selective plating using PM agar with 10 mM succinate but without (NH_4_)_2_SO_4_ or yeast extract for *R. palustris* and LB agar for *E. coli*. Statistical analyses were performed in GraphPad Prism 6.0.

## ACKNOWLEDGMENTS

We thank P.L. Foster for providing the KEIO collection, J. Liu at the IU Center for Genomics and Bioinformatics for assistance with Illumina sequencing, and members of the McKinlay laboratory for helpful discussions. This work was supported by U.S. Army Research Office grant no. W911NF-14-1-0411.

We declare no conflict of interest.

---

**Fig. S1.**
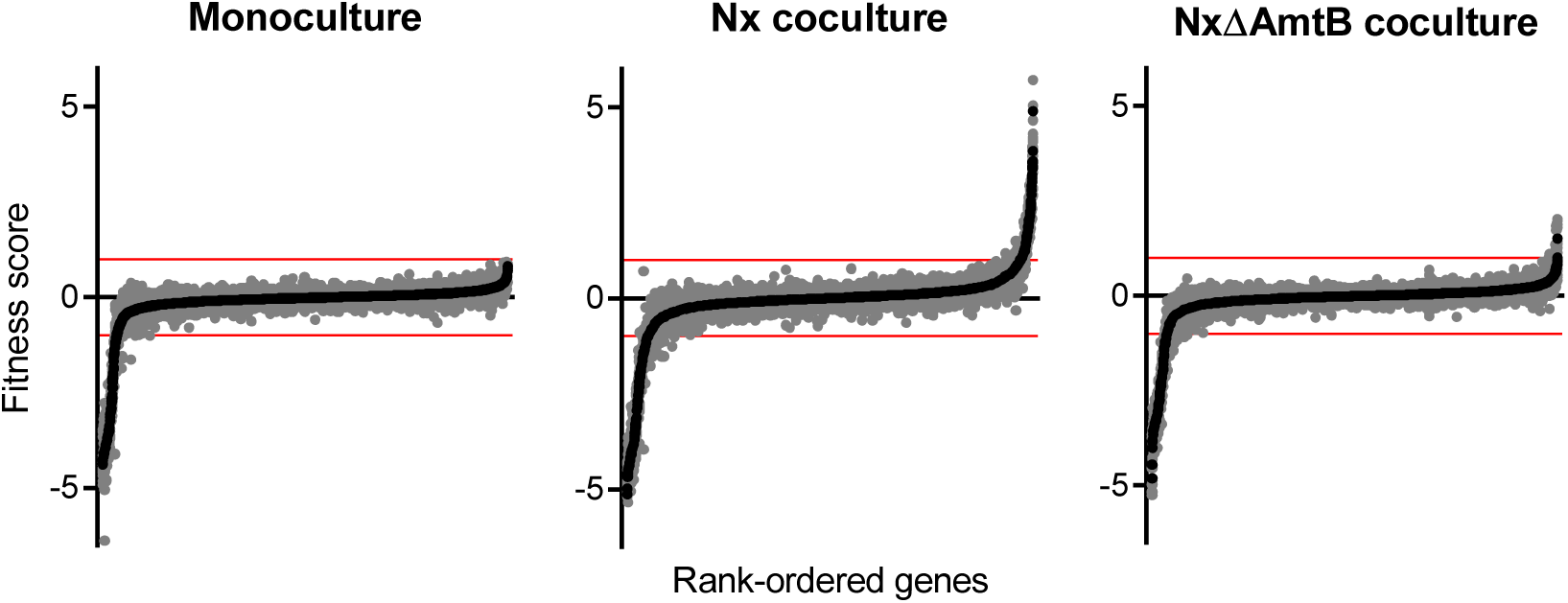
Rank-ordered mean fitness scores for 3,564 *E. coli* genes in monoculture or in cocultures with *R. palustris*. Genes were ranked from lowest to highest mean fitness score for each condition. Black circles indicate mean fitness scores, with gray circles indicating independent fitness scores for quadruplicate cultures. Red lines denote the threshold of |mean fitness|>1 used to identify strong fitness effects.

**Fig. S2.**
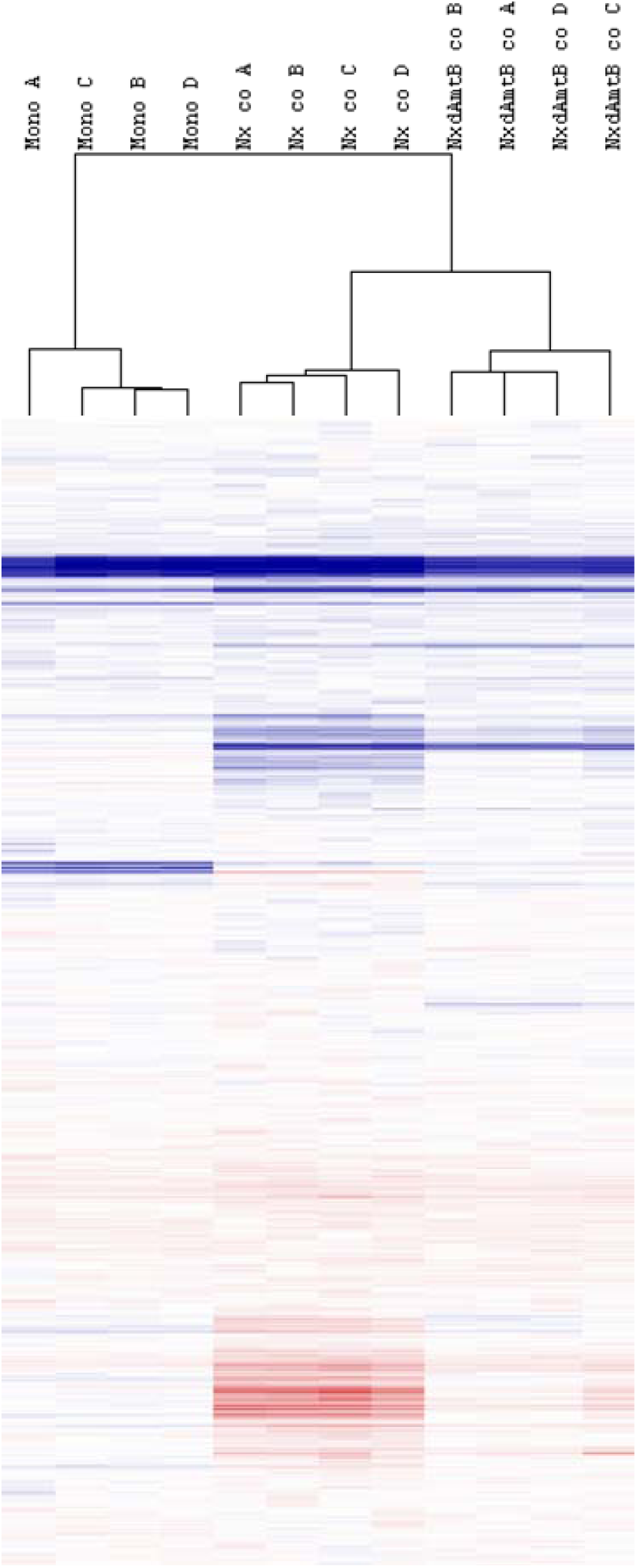
Dendrogram of fitness scores for all 3,564 *E. coli* genes for which fitness scores could be calculated. Genes and samples were clustered hierarchically (average linkage, uncentered correlation). Mono = monoculture; Nx co = Nx coculture; NxdAmtB co = NxΔAmtB coculture.

**Fig. S3.**
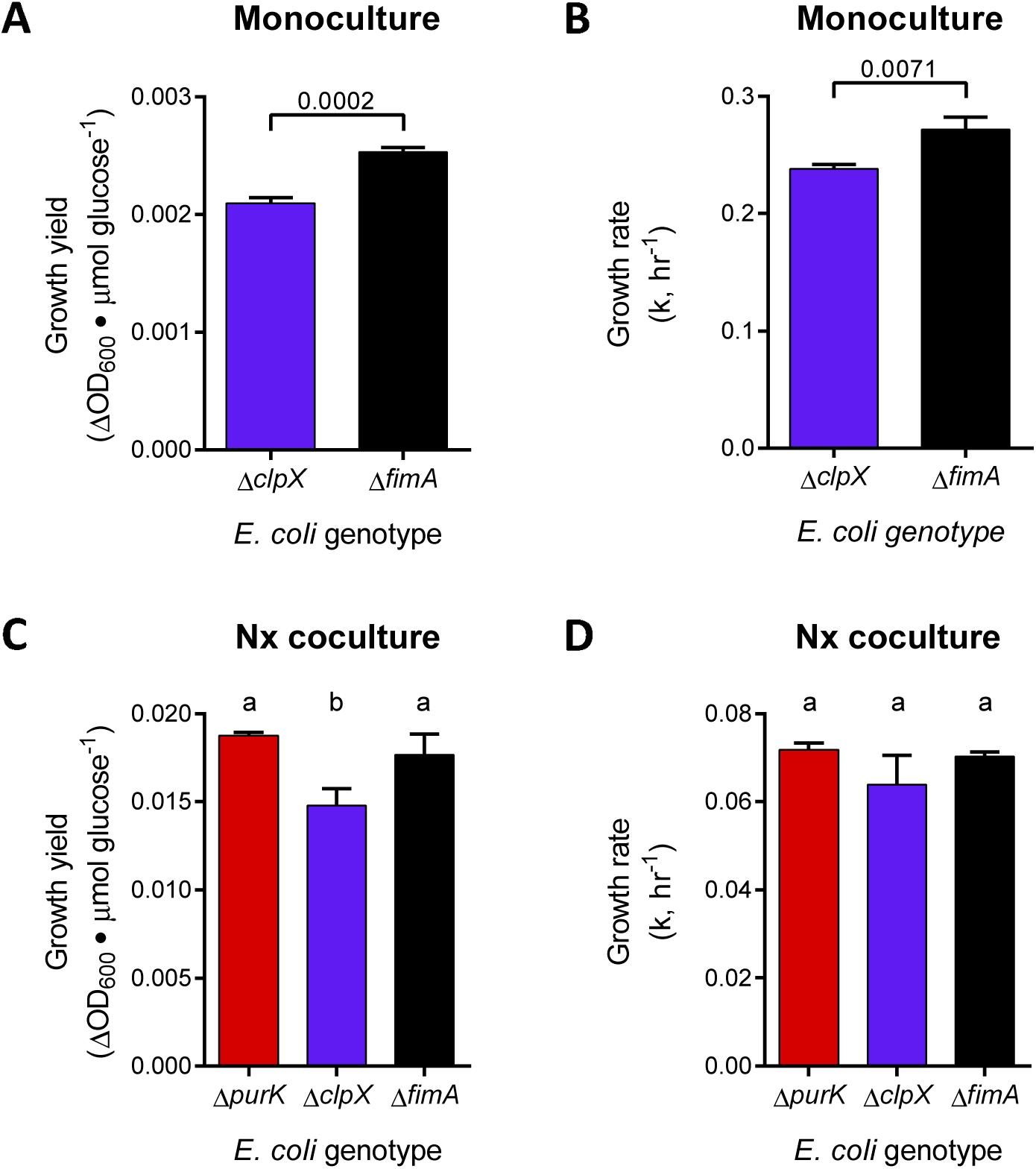
Growth yields (A, C) and exponential growth rates (B, D) for individual *E. coli* KEIO mutants grown in monoculture (A, B) or Nx coculture (C, D). Bars indicate the mean (±SD) of triplicate biological replicates. Final cell densities for determining growth yield were measured at the final time points in the respective growth curves in Fig. 3D, E. (A, B) Statistical differences were analyzed by unpaired t test and specific P values are indicated. (C, D) Different letters indicate statistical differences between groups, assessed by one-way analysis of variance with Tukey’s posttest (P<0.05).

